# DropSynth-Gold: Golden Gate Assembly in Emulsions Extends Multiplexed Gene Libraries to Greater Lengths

**DOI:** 10.64898/2026.05.29.728538

**Authors:** Karl J. Romanowicz, Samuel R. Hinton, Natanya Villegas, Calin Plesa

## Abstract

The ability to synthesize longer genes at scale remains a central challenge in multiplexed gene synthesis. DropSynth is a pooled gene synthesis platform that enables highly multiplexed, compartmentalized assembly from microarray-derived oligonucleotides, but current implementations rely on polymerase cycling assembly (PCA), which constrains fragment number, construct length, and assembly fidelity. Here we present DropSynth-Gold, an evolution of the DropSynth platform that replaces PCA with Golden Gate assembly (GGA) performed within emulsion droplets. This modification preserves the core workflow, including bead-linked oligonucleotide capture and pooled processing, while altering only the assembly chemistry and computational oligo design strategy. Emulsion-based Golden Gate assembly enables directional, multi-fragment ligation within isolated droplets, followed by recovery and amplification of full-length constructs. As a proof of concept, we constructed six 384-member libraries spanning increasing construct lengths and fragment counts, including designs from 5×300-mer fragments to 12×350-mer architectures (∼3 kb). DropSynth-Gold reliably assembled full-length constructs across all libraries. A direct comparison of a shared 5×300-mer library demonstrated comparable recovery and fidelity to PCA-based DropSynth, indicating that Golden Gate assembly can replace PCA without compromising assembly performance. These gains were achieved without increasing cost, workflow complexity, or turnaround time, expanding the accessible design space for multiplexed gene synthesis.

## Introduction

Multiplexed gene synthesis underpins many applications in synthetic biology, including functional genomics, protein engineering, and high-throughput screening (1-3). Advances in oligonucleotide synthesis have enabled the parallel generation of large DNA libraries with precisely defined sequences (4,5). However, because individual oligonucleotides are typically limited to ∼150-350 nucleotides, assembling longer gene constructs from pooled oligo libraries remains a central challenge (6,7). While several multiplexed assembly strategies have been developed (8,9), many lack effective compartmentalization, leading to cross-reactivity between constructs and limiting their ability to scale to longer genes or larger libraries (10).

To address these limitations, we previously developed DropSynth, a multiplexed gene synthesis method that enables pooled assembly of gene libraries within emulsion droplets (11). In this framework, oligonucleotides encoding each gene are captured onto uniquely barcoded microbeads and compartmentalized prior to assembly, ensuring that each droplet contains only the fragments required for a single construct (**Fig. 1A**). This architecture enabled scalable, low-cost assembly of large gene libraries and substantially improved upon prior multiplexed synthesis approaches (8,9,12). We subsequently extended the platform with DropSynth 2.0, improving assembly fidelity and library scale through optimization of polymerase choice, oligo design, and error correction strategies (13). However, both implementations rely on polymerase cycling assembly (PCA), in which overlapping oligonucleotides are joined through iterative polymerase extension. As construct length and fragment number increase, PCA becomes increasingly error-prone, generating recombination artifacts, reduced fidelity, and preferential amplification of shorter or partially assembled products (14-17). These limitations constrain reliable assembly at increasing fragment counts and construct lengths, making assembly chemistry a primary bottleneck despite advances in compartmentalization and scale (**Fig. 1A**).

**Figure 1.**
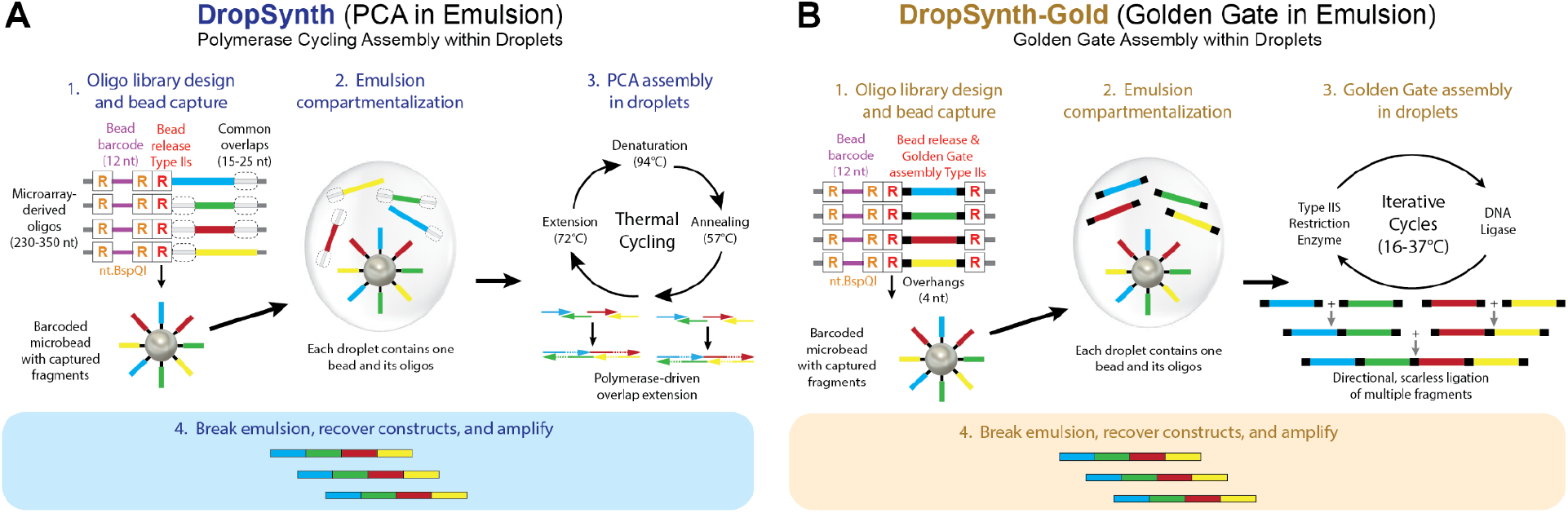
Assembly chemistry comparison. (**A**) DropSynth workflow using polymerase cycling assembly (PCA). Gene-specific oligonucleotides are captured on barcoded microbeads, compartmentalized within emulsion droplets, and assembled through iterative polymerase-driven extension, generating full-length constructs along with partial assemblies and chimeric byproducts. (**B**) DropSynth-Gold workflow using Golden Gate assembly (GGA). Oligonucleotide fragments containing Type IIS restriction sites and directional overhangs are captured on barcoded microbeads and assembled within droplets through iterative digestion-ligation cycles, enabling ordered multi-fragment assembly with reduced byproduct formation.

Recent work has demonstrated that Golden Gate assembly (GGA) can support highly multiplexed multi-fragment DNA assembly through computational selection of orthogonal Type IIS overhangs with high predicted ligation fidelity (18,19). Using these approaches, previous studies assembled constructs from dozens of fragments, including one-pot reconstruction of a 40 kb bacteriophage genome from 52 parts, as well as recovery of gene libraries from oligo pools using Golden Gate-compatible fragmentation and subpool amplification workflows (20-22). Unlike PCA-based assembly, which relies on extensive overlap hybridization between partially assembled intermediates, these methods use programmable ligation junctions to direct ordered fragment assembly and minimize recombination-driven byproducts. Together, these studies suggested that ligation-based assembly chemistries could provide a scalable alternative for highly multiplexed gene synthesis.

Here, we present DropSynth-Gold, which addresses the limitations of PCA by replacing it with Type IIS restriction enzyme-mediated GGA within droplets (**Fig. 1B**). This modification preserves the core DropSynth workflow, including multiplexed bead-based oligo capture, emulsion compartmentalization, and pooled recovery, while fundamentally changing the assembly mechanism. In contrast to polymerase-driven overlap extension, GGA uses programmable overhangs to direct ordered ligation of DNA fragments through iterative digestion-ligation cycles (23-25), reducing recombination artifacts and improving scalability with increasing fragment number (**Fig. 1B**).

To evaluate this approach, we constructed six 384-member gene libraries spanning increasing construct lengths and fragment counts, from 5×300-mer assemblies to 12×350-mer architectures approaching 3 kb total construct length. We performed a direct head-to-head comparison of PCA-based DropSynth and GGA-based DropSynth-Gold using a shared 5×300-mer library, demonstrating comparable recovery and fidelity between assembly strategies. We additionally developed a computational framework to redesign oligonucleotide architectures for Golden Gate compatibility, incorporating directional overhang selection, restriction site filtering, and fragment partitioning constraints without increasing experimental burden. Together, these results establish DropSynth-Gold as an alternative assembly strategy that maintains performance comparable to PCA while extending multiplexed gene assembly to longer constructs composed of more fragments.

## Materials and Methods

### Oligonucleotide Design for DropSynth and DropSynth-Gold

Oligonucleotide design for PCA-based DropSynth libraries was performed as previously described (11,13), including codon optimization, restriction site filtering, and partitioning of gene sequences into overlapping oligos constrained by melting temperature and secondary structure. Briefly, amino acid sequences derived from either FPBase (26) or the reference UniProt human proteome (27) were length filtered and reverse translated using *E. coli*-optimized codon frequencies with DNA Chisel (28). Coding sequences were flanked with cloning and amplification sequences and split into overlapping oligos for polymerase cycling assembly.

For DropSynth-Gold, oligonucleotide design was modified to support Golden Gate assembly. The same amino acid inputs were used, but coding sequences were partitioned into discrete 300–350 nt fragments flanked by BsaI recognition sites and directional 4-bp overhangs. Gene termini were additionally designed with 4-nt poly(A) overhangs to enable release of fragments from bead-bound oligonucleotides while preventing participation of gene ends in GGA ligation reactions. Fragment boundaries were selected so junction overhangs corresponded to the endogenous sequence, enabling scarless assembly (**Fig. S1**). Candidate junctions were constrained by fragment size, oligo payload length, and positional tolerance relative to equal-sized splits. Overhangs were filtered to exclude homopolymers and palindromes, maintain balanced GC content, enforce minimum sequence distance, and minimize predicted mis-ligation using the Potapov et al. (29) ligation fidelity dataset through Tatapov (available from https://github.com/Edinburgh-Genome-Foundry/tatapov). Candidate overhangs were required to exceed a minimum self-annealing threshold and remain below a maximum cross-annealing threshold based on the 37 °C, 1 h Potapov dataset (29). Internal Type IIS sites were removed by codon reassignment prior to fragment partitioning. Final fragment sets were validated computationally by confirming overhang compatibility and accurate reconstruction of the original coding sequence. Scripts were implemented in Python using Biopython and Tatapov, with optional DNA Cauldron-based Golden Gate simulation for selected designs.

### Oligo Synthesis, Amplification, and Bead Loading

Oligonucleotide pools were synthesized by Twist Bioscience (San Francisco, CA, USA). Oligo amplification, processing, and bead loading were performed as previously described for DropSynth 2.0 (13). Briefly, oligonucleotide subpools were PCR-amplified using KAPA HiFi DNA Polymerase (Roche), with qPCR used to determine cycle numbers and minimize overamplification bias, followed by silica column purification (GeneJET, Thermo Fisher Scientific). Oligos were then processed by nt.BspQI nicking to expose 12-nt barcode overhangs, loaded onto complementary DNA-barcoded microbeads as previously described (11,13), and purified by streptavidin bead capture and column cleanup. Processed oligos were annealed and ligated to ∼5 × 10^6^ barcoded microbeads using a slow temperature ramp to promote specific barcode hybridization and co-localization of all oligos for a given construct on a single bead. Excess unbound oligos were removed by sonication and washing prior to emulsion assembly.

### PCA-Based Emulsion Assembly in Droplets

For the PCA-based DropSynth control library, oligo-loaded beads were emulsified and assembled using polymerase cycling assembly as previously described (11,13). Briefly, bead-containing reaction mixtures were combined with EvaGreen (Bio-Rad) droplet generation oil and emulsified by vortexing (∼3,000 rpm, 3 min; Vortex Genie 2), compartmentalizing individual beads within micron-scale droplets. Reactions were then incubated at 37 °C to release oligonucleotides from beads via BtsI-v2 cleavage (New England Biolabs), followed by thermocycling to enable overlap annealing and polymerase extension into full-length products. Reaction conditions were optimized to maximize full-length assembly while minimizing incomplete products and recombination artifacts.

### Golden Gate-Based Emulsion Assembly in Droplets

For DropSynth-Gold, assembly reactions were modified to incorporate GGA within droplets. Bead-loaded oligos were combined with EvaGreen droplet generation oil and emulsified by vortexing (∼3,000 rpm, 3 min) to generate compartmentalized droplets. Assembly reactions contained BsaI-HFv2 (New England Biolabs), T4 DNA ligase, 10x T4 DNA ligase buffer supplemented 1:1 with ATP, recombinant bovine serum albumin (rBSA), and nuclease-free water. Emulsified reactions were subjected to 200 thermocycles alternating between 37 °C (digestion) and 16 °C (ligation) to enable iterative restriction-ligation assembly of fragments within droplets. Following cycling, reactions were incubated at 60 °C for 5 min to inactivate T4 DNA ligase and at 80 °C for 20 min to inactivate BsaI-HFv2.

### Emulsion Breaking and DNA Recovery

Emulsions were broken using chloroform-based phase separation for both PCA-based DropSynth and DropSynth-Gold assemblies. Following assembly, reactions were split into 1.5 mL microcentrifuge tubes (∼350 µL per tube) where 175 uL of nuclease-free water and 615 µL of chloroform was added to each tube, and samples were vortexed at maximum speed (∼3,000 rpm; Vortex Genie 2) for 3 min to disrupt the emulsion. Tubes were sealed with high vacuum grease and centrifuged at 16,000 RCF for 10 min at room temperature to achieve phase separation. The aqueous phase was carefully recovered and purified using a GeneJET column-based PCR cleanup kit.

### Library Recovery and Amplification

Recovered DNA was separated by electrophoresis on a 2% agarose gel (110 V, 35 min). Bands corresponding to the expected assembly sizes were excised and purified by gel extraction and subsequent column cleanup. For the PCA-based DropSynth library, the size-selected product was amplified using a single suppression PCR primer (sPCR) to enrich for full-length assemblies, as previously described (13). Suppression PCR leverages inverted terminal repeat (ITR) sequences flanking assembled products, which suppress amplification of shorter fragments through intramolecular annealing. Reactions were performed using KAPA HiFi DNA Polymerase, with cycle numbers determined by qPCR and amplification stopped in the late exponential phase.

For DropSynth-Gold libraries, size-selected products were amplified using standard PCR with KAPA HiFi DNA Polymerase. Amplification cycles were determined by qPCR and optimized to preserve library representation and minimize overamplification. Amplified libraries were purified by column cleanup and quantified prior to sequencing.

### Sequencing and Analysis

Recovered libraries were amplified using primers containing PacBio Kinnex compatible adapters and library-specific flanking sequences to enable multiplexed long-read sequencing. Amplified products were purified, quantified, and submitted to the University of Oregon Genomics and Cell Characterization Core Facility (GC3F) for Kinnex array generation and PacBio Revio sequencing. PacBio CCS reads were deconcatenated with Skera and demultiplexed with Lima to generate high-quality consensus reads, and downstream analysis was performed using reproducible Python and R workflows. FASTA files for each library were processed independently to compute summary statistics, including read counts, perfect read fraction, and coverage metrics. Resulting summary statistics were subsequently aggregated across libraries for downstream analysis and visualization. Reads were filtered at two quality thresholds depending on the analysis. Sequence fidelity was assessed using stringent Q40-filtered reads to ensure high-confidence base calls, whereas library coverage and uniformity were evaluated using Q30-filtered reads to retain greater sequencing depth while maintaining high read quality. At both filtering levels, reads were further restricted to those within ±20 bp of the expected reference length to remove truncated and off-target products. Filtered reads were aligned to the corresponding reference library sequences using minimap2 (30), retaining only primary alignments. Mapped reads were parsed to assign each read to its corresponding reference construct. Library coverage was defined as the number of unique reference sequences represented by at least one mapped read. Sequence fidelity was assessed at the DNA level using Q40-filtered reads. Reads were classified as perfect assemblies if they matched the expected reference sequence exactly across the full-length construct. The perfect read fraction (*P*) was calculated as the proportion of mapped Q40-filtered reads classified as perfect assemblies. Per-base error rates (reported as 1:X bp) were then estimated as 1/(1−*P*^1/*L*^), where *L* is construct length. For Q30-filtered reads, additional metrics were computed to assess library-level performance, including coverage, representation uniformity, and unique perfect sequence recovery. Additional summary statistics, including read length distributions, alignment identity, and indel frequency, were calculated to characterize overall assembly quality. All analysis parameters were held constant across libraries to enable direct comparisons.

### Library Construction and Benchmarking

To evaluate performance across increasing construct lengths and fragment counts, we constructed six gene libraries (L1-L6) of 384 members, except L3 (270 members), spanning assembly architectures from 5 to 12 fragments and approximately 1.0 kb to 3.0 kb in total construct length. Libraries were designed using either 300-mer or 350-mer oligonucleotides, including 5×300-mer (L1-L2), 8×300-mer (L3), and larger assemblies composed of 350-mer fragments (L4-L6) (**Table 1**).

**Table 1.**
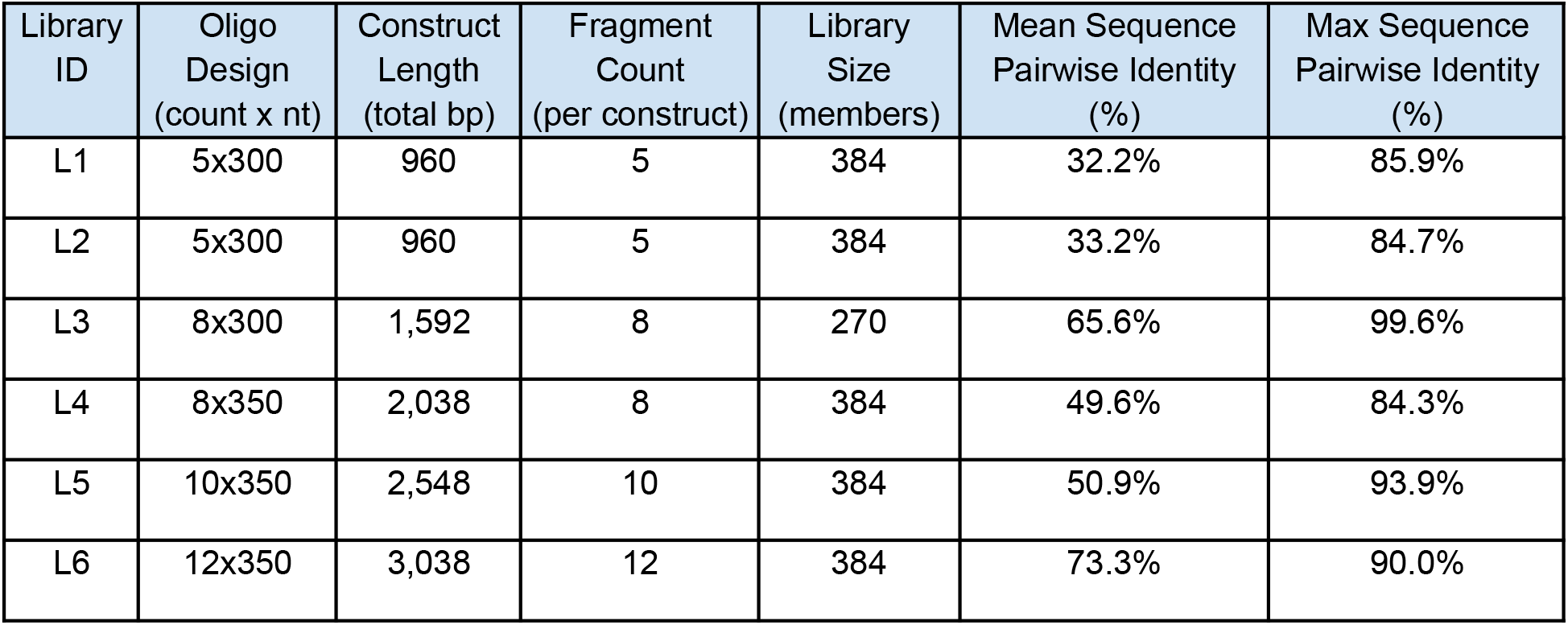
Design of benchmark libraries spanning increasing construct lengths and fragment numbers. Six gene libraries (L1-L6) were constructed to benchmark DropSynth-Gold across increasing fragment numbers and construct lengths, spanning assembly architectures from 5 to 12 fragments and approximately 1.0 kb to 3.0 kb. Libraries were designed using either 300-mer or 350-mer oligonucleotides. Sequence designs were derived from FPBase or the UniProt human proteome and filtered by length to match target construct sizes. Mean and maximum pairwise sequence identity were calculated for each library to quantify sequence diversity. Amino acid sequences served as inputs for codon optimization and Golden Gate-compatible oligonucleotide design workflows for DropSynth-Gold assembly. Library size was held constant across conditions except for L3, which contained 270 members due to design constraints.

| Library ID | Oligo Design (count x nt) | Construct Length (total bp) | Fragment Count (per construct) | Library Size (members) | Mean Sequence Pairwise Identity (%) | Max Sequence Pairwise Identity (%) |
| --- | --- | --- | --- | --- | --- | --- |
| L1 | 5x300 | 960 | 5 | 384 | 32.2% | 85.9% |
| L2 | 5x300 | 960 | 5 | 384 | 33.2% | 84.7% |
| L3 | 8x300 | 1,592 | 8 | 270 | 65.6% | 99.6% |
| L4 | 8x350 | 2,038 | 8 | 384 | 49.6% | 84.3% |
| L5 | 10x350 | 2,548 | 10 | 384 | 50.9% | 93.9% |
| L6 | 12x350 | 3,038 | 12 | 384 | 73.3% | 90.0% |

Libraries were designed to systematically increase fragment number and construct length while maintaining a consistent library size (384 members; L3, 270 members), isolating construct length and fragment count as the primary variables (**Table 1**). To assess sequence diversity within each library, mean and maximum pairwise sequence identity were calculated across all protein designs. Following oligo design, pooled oligonucleotide libraries were synthesized, amplified, assembled, size selected, sequenced, and processed as described above.

## Results

### Golden Gate Enables High-Fidelity Gene Assembly within Emulsion Droplets

To establish Golden Gate assembly within the DropSynth framework, we evaluated whether Type IIS restriction-ligation chemistry could support multi-fragment gene assembly within emulsion droplets. Bead-bound oligonucleotide libraries were compartmentalized and subjected to iterative digestion-ligation cycling using BsaI-HFv2 and T4 DNA ligase (**Fig. 1B**).

As an initial proof-of-concept, we directly compared conventional PCA-based DropSynth and GGA-based DropSynth-Gold using the same L1 library (5×300-mer design) (**Fig. S2**). Sequencing analysis demonstrated that GGA achieved performance comparable to PCA while supporting efficient recovery of correctly assembled products. Notably, DropSynth-Gold produced a dominant full-length assembly peak with only minor discrete byproduct peaks corresponding to partial 2-, 3-, and 4-fragment assemblies (**Fig. S3**). This pattern suggests that GGA generates more well-defined intermediates that should facilitate more effective size selection compared to the broader smeared byproducts commonly observed in PCA-based assemblies. The distribution of per-gene perfect assemblies, defined as Q30-filtered reads matching the reference sequence exactly at the DNA level (NM = 0), was broadly similar between workflows (**Fig. 2A**). Error rates estimated from Q40-filtered reads were also comparable, improving from approximately 1:951 bp for PCA-based DropSynth to 1:1196 bp for DropSynth-Gold.

**Figure 2.**
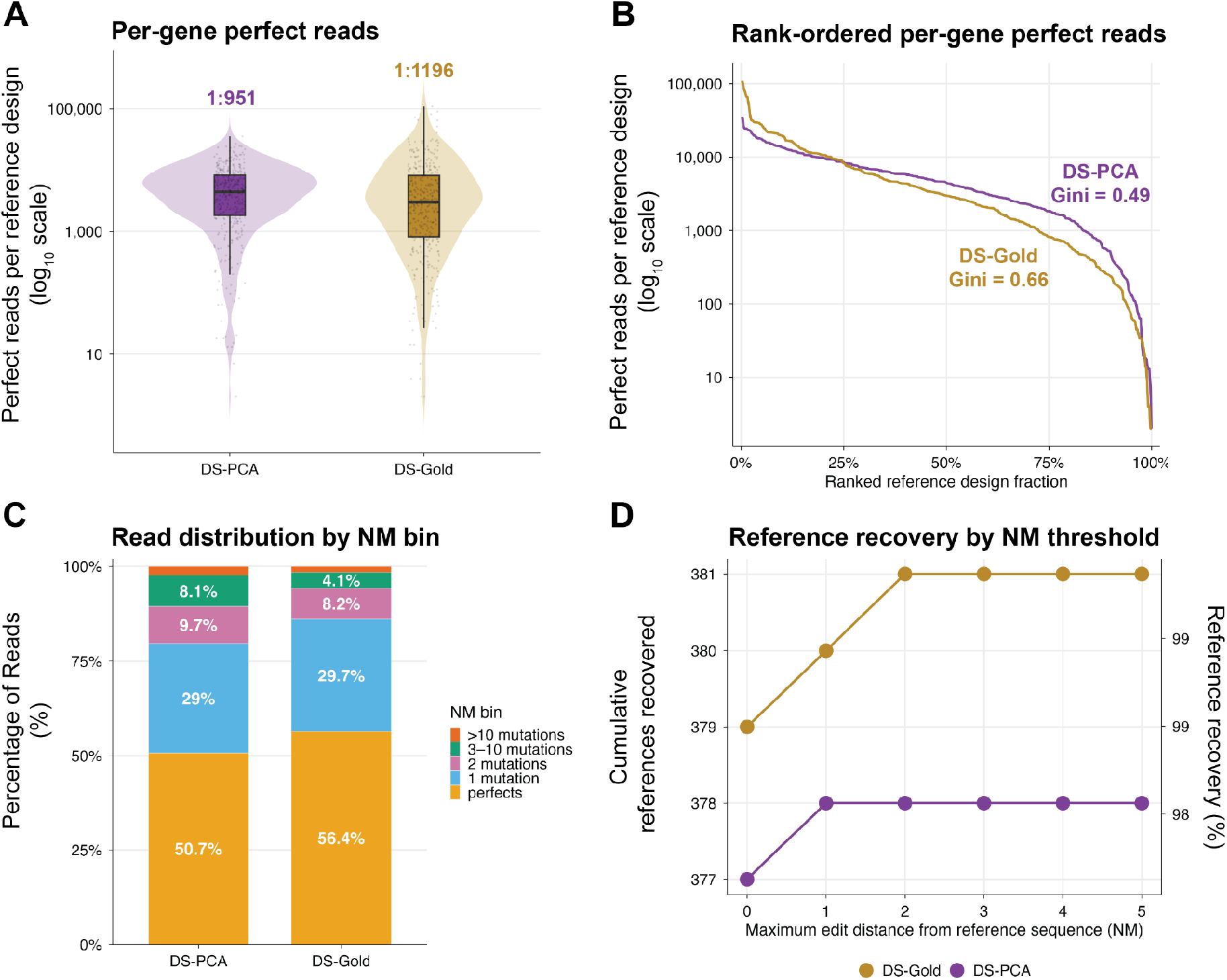
DropSynth-Gold supports multiplexed gene assembly with comparable library recovery relative to PCA-based DropSynth. (**A**) Distribution of per-gene perfect read counts for the L1 library (5×300-mer design) comparing PCA-based DropSynth and DropSynth-Gold. Per-gene perfect reads were calculated from Q30-filtered sequencing data and defined as reads matching the reference sequence exactly at the DNA level (NM = 0). Violin plots show distributions across reference designs, with boxplots indicating median and interquartile range. Estimated assembly error rates calculated from Q40-filtered reads are overlaid above each distribution. (**B**) Rank-ordered distributions of per-gene perfect read counts across the L1 library. Gini coefficients are shown for each assembly workflow, where lower values indicate more even representation across designs. (**C**) Distribution of sequencing reads across edit distance (NM) bins for PCA-based DropSynth and DropSynth-Gold libraries. Reads were filtered to retain sequences within ±20 bp of expected reference length prior to alignment analysis. (**D**) Coverage and cumulative reference recovery as increasingly permissive edit-distance thresholds were applied (NM ≤ 0 through NM ≤ 5).

Rank-ordered abundance analysis further demonstrated broad library recovery for both assembly strategies (**Fig. 2B**). PCA-based DropSynth and DropSynth-Gold each achieved >98% library coverage, although PCA-based assembly exhibited greater representation uniformity across constructs. This difference was reflected by a lower Gini coefficient for PCA-based DropSynth (0.49) relative to DropSynth-Gold (0.66), where lower values indicate more uniform representation across library members.

To further characterize assembly accuracy, we quantified read distributions across edit-distance (NM) bins following filtering to retain reads within ±20 bp of the expected reference length, reducing contributions from truncated products and large insertion/deletion events (**Fig. 2C**). Perfect assemblies (NM = 0) represented the largest read fraction in both workflows, accounting for 50.7% of reads in PCA-based DropSynth and 56.4% of reads in DropSynth-Gold. An additional substantial fraction of reads differed by only one mutation (29.0% and 29.7%, respectively), while progressively fewer reads were observed at higher edit distances.

Consistent with this observation, coverage and cumulative reference recovery saturated rapidly as increasingly permissive edit-distance thresholds were applied (**Fig. 2D**). PCA-based DropSynth achieved 98.2% coverage (377/384 reference designs) using only perfect reads and plateaued at 98.4% coverage (378/384) by NM ≤ 1, whereas DropSynth-Gold achieved 98.7% coverage (379/384) using perfect reads and reached 99.2% coverage (381/384) by NM ≤ 2. Minimal gains beyond these thresholds indicate that recovered library members were predominantly represented by exact or near-exact sequence matches rather than highly mutated assemblies.

### DropSynth-Gold Supports Longer Constructs and Higher Fragment Counts

To evaluate the scalability of DropSynth-Gold, we constructed six multiplexed libraries ranging from ∼1.0-3.0 kb and assembled from 5-12 fragments (L1-L6; **Table 1**). Libraries L1, L2, L4, L5, and L6 each contained 384 reference designs, while L3 contained 270 designs due to sequence constraints. Together, these libraries systematically increased construct length and fragment number while maintaining a consistent pooled assembly workflow, enabling evaluation of assembly performance across a broad range of library sizes and architectures (**Fig. S2**). Previous attempts to extend PCA-based DropSynth assemblies beyond five 300-mer fragments produced poor assembly outcomes (data not shown), likely due to increased cross-hybridization and misassembly among large numbers of overlapping oligonucleotides.

Sequencing analysis demonstrated that DropSynth-Gold supports accurate multiplexed assembly across increasing construct lengths and fragment counts (**Fig. 3A**). Per-gene perfect assemblies generally decreased with increasing construct length and fragment number. Shorter libraries (L1 and L2; 5×300-mer assemblies, ∼1 kb) exhibited the highest abundance of perfect reads per design and Q40-derived error rates of approximately 1:1196 bp and 1:1307 bp, respectively. Libraries L3 and L4 (∼1.6-2.0 kb) maintained comparable error rates of approximately 1:1185 bp and 1:1391 bp, while larger assemblies L5 (10×350; ∼2.5 kb) and L6 (12×350; ∼3.0 kb) exhibited error rates of approximately 1:1212 bp and 1:1141 bp. Overall, assembly fidelity remained broadly consistent across all libraries evaluated.

**Figure 3.**
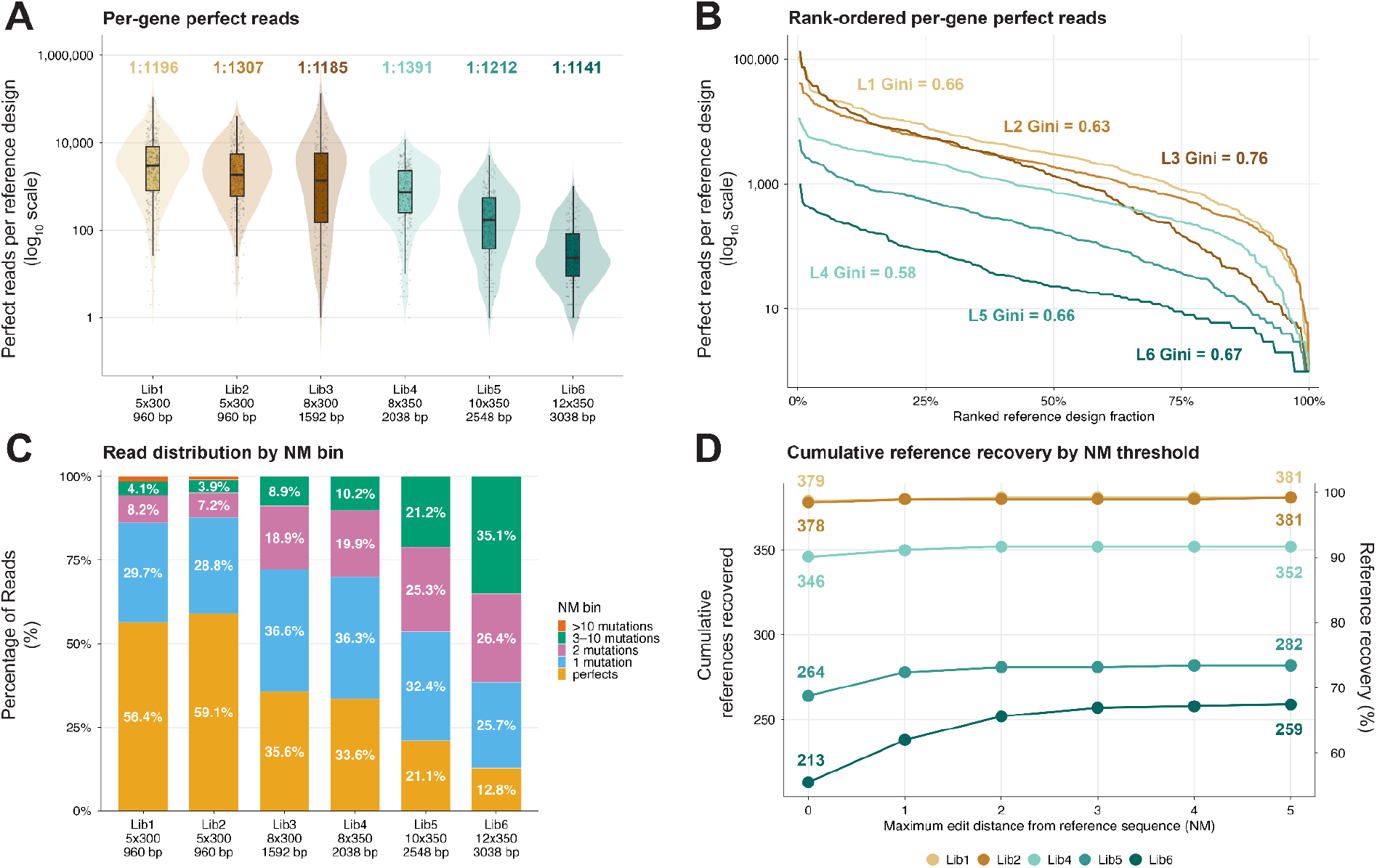
DropSynth-Gold supports multiplexed gene assembly across increasing fragment number and construct length. (**A**) Distribution of per-gene perfect read counts across DropSynth-Gold libraries L1-L6 spanning 5-12 fragments and construct lengths from ∼1.0 kb to ∼3.0 kb. Per-gene perfect reads were calculated from Q30-filtered sequencing data and defined as reads matching the reference sequence exactly at the DNA level (NM = 0). Violin plots show distributions across reference designs, with boxplots indicating median and interquartile range. Estimated assembly error rates from Q40-filtered reads are overlaid above each distribution. (**B**) Rank-ordered distributions of per-gene perfect read counts across libraries L1-L6. Gini coefficients are shown for each library, where lower values indicate more even representation across designs. (**C**) Distribution of sequencing reads across edit distance (NM) bins for libraries L1-L6. Reads were filtered to retain sequences within ±20 bp of expected reference length prior to alignment analysis. (**D**) Coverage and cumulative reference recovery as increasingly permissive edit-distance thresholds were applied (NM ≤ 0 through NM ≤ 5). Coverage was calculated as the fraction of reference designs recovered at each threshold. Library L3 was excluded because its reduced library size (270 reference designs versus 384 for all other libraries) precluded direct comparison of recovery percentages.

Rank-ordered abundance analysis further demonstrated robust recovery across all libraries (**Fig. 3B**). Shorter assemblies generally exhibited higher per-design abundance and greater representation uniformity, whereas longer, higher-fragment assemblies showed greater representation skew across pooled constructs. Libraries L1, L2, and L4 maintained comparatively even representation (Gini coefficients 0.66, 0.63, and 0.58, respectively), whereas L3 exhibited the greatest skew (Gini = 0.76). The longest libraries, L5 and L6, also showed elevated skew (Gini coefficients 0.66 and 0.67, respectively), suggesting that increasing construct length and fragment count contribute to representation bias during pooled assembly and recovery. The elevated Gini coefficient observed for L3 may additionally reflect use of a partial barcode set (270 reference designs versus 384 for all other libraries), potentially increasing susceptibility to uneven pooled representation or incomplete construct membership.

Perfect assemblies (NM = 0) represented the dominant read class in shorter libraries (**Fig. 3C**), accounting for 56.4% and 59.1% of reads in L1 and L2, respectively. Increasing construct length and fragment count shifted read distributions toward low-edit-distance variants (NM = 1-10), whereas high-edit-distance categories remained comparatively infrequent (NM > 10). Libraries L3 and L4 retained substantial fractions of perfect assemblies, whereas the longest libraries showed the greatest shift toward near-perfect sequence variants. For example, L6 retained 12.8% perfect reads while maintaining substantial representation within low-edit-distance categories, indicating that longer, higher-fragment assemblies primarily introduced modest sequence deviations rather than highly mutated products.

Coverage increased rapidly as increasingly permissive edit-distance thresholds were applied (**Fig. 3D**). Shorter libraries achieved near-complete recovery using only perfect reads, with L1 recovering 379 of 384 reference designs and L2 recovering 381 of 384 designs at NM = 0. Longer libraries exhibited lower exact-match recovery but substantial gains when low-edit-distance reads were included. For example, L5 increased from 264 recovered reference designs at NM = 0 to 282 at NM ≤ 5, representing a 6.8% increase in recovered designs, while L6 improved from 213 to 259 recovered reference designs, a 21.6% increase. Reduced recovery of unique reference designs in L5 and L6 was likely influenced in part by lower sequencing depth, with L5 and L6 yielding 2,890,304 and 870,788 usable reads, respectively, compared to approximately 6-12 million usable reads for L1-L4. Minimal gains beyond NM ≤ 2-5 indicate that recovery losses in longer, higher-fragment libraries were driven primarily by near-perfect sequence variants rather than highly mutated assemblies. L3 was excluded from cumulative recovery comparisons because its reduced library size (270 reference designs) precluded direct percentage-based comparison with the 384-member libraries.

### Assembly Fidelity Remains Stable Across Increasing Lengths and Fragments

To further evaluate how assembly performance scales across increasing construct lengths and fragment numbers, we examined estimated assembly error rates across DropSynth-Gold libraries spanning ∼1.0-3.0 kb constructs assembled from 5-12 fragments (**Fig. 4**). Estimated assembly fidelity remained broadly consistent across libraries, with no evidence of abrupt performance deterioration at longer construct lengths or higher fragment counts. Q40-filtered error estimates ranged from approximately 1:1141 bp to 1:1391 bp across libraries L1-L6, while Q30-filtered estimates ranged from approximately 1:685 bp to 1:774 bp. Intermediate-length assemblies (∼2.0 kb; L4) exhibited the highest estimated fidelity, whereas constructs approaching ∼3 kb maintained error rates comparable to shorter assemblies. Transitioning from 300-mer oligonucleotide designs (L1-L3) to 350-mer architectures (L4-L6) did not measurably reduce assembly fidelity despite increasing construct length and larger oligonucleotide building blocks. Libraries spanning 8×350-, 10×350-, and 12×350-mer assemblies maintained estimated Q40 error rates between approximately 1:1141 bp and 1:1391 bp. Although modest library-to-library variability was observed, assembly performance remained stable across the construct range evaluated, indicating that GGA scales predictably within the DropSynth framework without substantial degradation in sequence accuracy.

**Figure 4.**
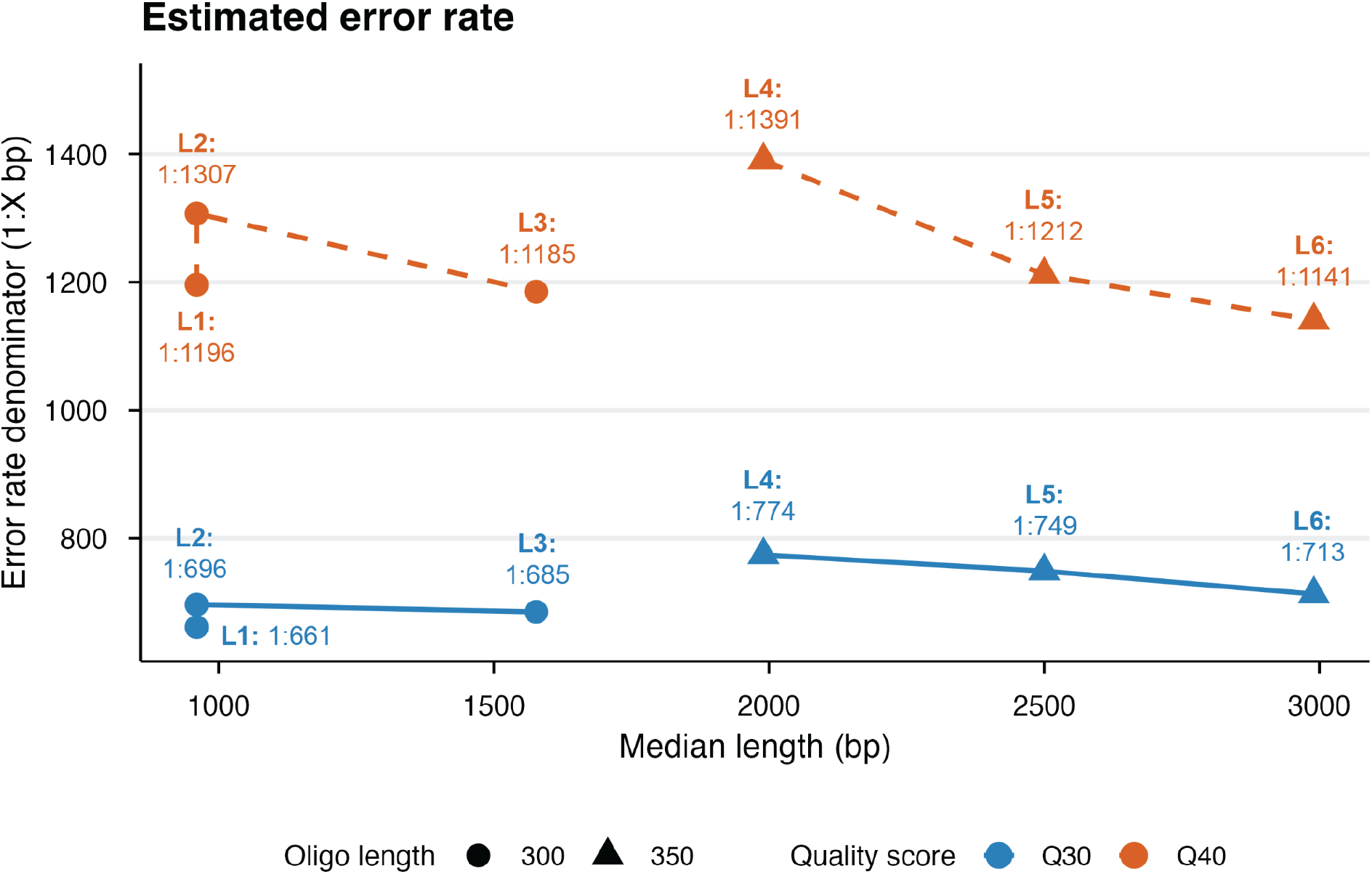
Assembly fidelity across increasing construct lengths and fragment numbers. Estimated assembly error rates across DropSynth-Gold libraries L1-L6 spanning construct lengths from ∼1.0-3.0 kb and assemblies composed of 5-12 oligonucleotide fragments. Error rates were estimated from Q30- and Q40-filtered sequencing reads and are shown as denominator values (1:X bp). Circular markers indicate 300-mer oligonucleotide architectures (L1-L3), whereas triangular markers indicate 350-mer architectures (L4-L6).

## Discussion

DropSynth-Gold establishes Golden Gate assembly (GGA) within the DropSynth framework (11,13) as an effective strategy for multiplexed pooled gene synthesis. Direct comparison against PCA-based DropSynth demonstrated that GGA performs efficiently within emulsion droplets while maintaining comparable library recovery, construct representation, and assembly fidelity (**Fig. 2**). These results establish that Type IIS restriction-ligation chemistry can function robustly under droplet compartmentalization and provide an alternative assembly strategy for pooled gene synthesis without compromising overall library performance.

Benchmarking across construct lengths spanning ∼1-3 kb and assembly architectures ranging from five to twelve fragments demonstrated that DropSynth-Gold scales predictably across increasing construct lengths and fragment numbers (**Fig. 3**). Although exact assembly recovery declined for longer, higher-fragment libraries, assembly fidelity remained broadly stable, with Q40-derived error estimates remaining relatively consistent across libraries. Importantly, recovery improvements achieved through permissive edit-distance thresholds showed that longer assemblies primarily accumulated near-perfect sequence variants rather than highly mutated products. Constructs approaching ∼3 kb retained substantial library coverage and reference recovery without evidence of abrupt fidelity loss, indicating that GGA remains effective across substantially larger assembly architectures than previously demonstrated within PCA-based DropSynth workflows (11,13,31-33).

Representation uniformity remained broadly preserved across pooled libraries, although longer assemblies generally exhibited greater representation skew. This trend likely reflects a combination of reduced exact recovery, uneven amplification, and assembly-associated biases that become more pronounced with increasing construct length and fragment number. Unlike PCA-based assembly, GGA lacks an assembly-phase amplification step, meaning final construct yield is constrained by the least abundant fragment in the reaction. Consequently, GGA may be more sensitive to input oligo pool uniformity, as underrepresented fragments can directly limit full-length assembly recovery. This effect may contribute to the higher Gini coefficients observed in DropSynth-Gold relative to PCA-based DropSynth.

Notably, representation skew did not scale strictly with construct length. Library L3 exhibited the greatest skew despite its intermediate construct length, suggesting that factors beyond assembly architecture may also influence recovery behavior. Unlike the other benchmarking libraries, which each contained 384 reference designs, L3 contained only 270 members due to design constraints. L3 also exhibited the highest maximum pairwise sequence identity (99.6%) and elevated mean pairwise identity (65.6%) relative to most other libraries (**Table 1**), potentially increasing susceptibility to cross-ligation, uneven representation, or incomplete recovery during pooled assembly. Together, these observations suggest that sequence composition, library diversity, and uneven barcoded bead coverage can influence pooled recovery alongside assembly chemistry and construct architecture.

Transitioning from 300-mer to 350-mer oligonucleotide architectures did not substantially alter assembly fidelity despite increasing construct length and fragment number (**Fig. 4**). This observation suggests that oligonucleotide architecture itself was not a dominant driver of performance within the ranges evaluated here and further supports the robustness of GGA across diverse assembly designs. Nevertheless, longer assemblies exhibited reduced exact assembly recovery and modest increases in representation skew, highlighting opportunities for further optimization of larger libraries. Improvements in oligonucleotide synthesis quality, fragment design constraints, barcode recovery efficiency, and reaction conditions may further improve pooled assembly performance. Because input oligonucleotide quality remains a major determinant of final construct fidelity, advances in upstream synthesis quality will likely translate directly into downstream assembly improvements.

Compared to other Golden Gate-based multiplexed gene synthesis methods, DropSynth-Gold achieves substantially greater library scale by performing each assembly within an isolated emulsion droplet. This compartmentalized architecture allows library size to be determined primarily by barcode diversity rather than by the number of orthogonal GGA junctions that can coexist within a shared reaction. For example, OMEGA (22) can theoretically assemble up to 576 distinct 2.6 kb constructs per 96-well plate under its highest multiplexing regime, but scaling remains constrained by the finite set of orthogonal GGA junctions available within each subpool. In contrast, DropSynth-Gold retains the barcoded bead-in-emulsion architecture of DropSynth, enabling all assemblies to be performed simultaneously within a single pooled reaction using highly multiplexed bead sets containing 384, 1,536, or 12,288 unique barcodes (*preprint forthcoming*). Consequently, library scale can be expanded through barcode multiplexing without requiring hundreds of independently amplified and assembled subpools.

Other multiplexed DNA assembly approaches face similar throughput limitations. Sidewinder (34), which uses DNA three-way junctions to assemble fragments from oligonucleotide pools, reported a maximum pool size of only 24 distinct constructs (35). In addition, Sidewinder requires separate synthesis of both the top and bottom strands for each assembled fragment, effectively doubling oligonucleotide synthesis requirements relative to DropSynth-Gold. Together, these comparisons highlight how compartmentalized assembly enables DropSynth-Gold to achieve substantially greater throughput while maintaining the scalability advantages of pooled gene synthesis.

Collectively, these results extend the capabilities of pooled multiplexed gene synthesis within the DropSynth framework. By integrating Golden Gate assembly into a compartmentalized bead-in-emulsion workflow, DropSynth-Gold supports assembly architectures spanning up to twelve fragments and approximately 3 kb construct lengths while maintaining stable assembly fidelity, broad library recovery, and substantial representation of both exact and near-exact assemblies. These capabilities enable access to longer proteins, multi-domain constructs, and increasingly diverse design spaces that remain difficult to generate using existing pooled assembly methods. As oligonucleotide synthesis quality, barcode multiplexing capacity, and assembly chemistries continue to advance, DropSynth-Gold provides a scalable foundation for increasingly large and sophisticated gene libraries spanning applications in protein engineering, functional genomics, synthetic biology, and large-scale biological design.

## Acknowledgements

This work was supported in part by a grant from the Chan Zuckerberg Initiative DAF, an advised fund of Silicon Valley Community Foundation. We thank the University of Oregon Genomics and Cell Characterization Core Facility (GC3F) staff for assistance with next-generation sequencing. We are also grateful to Craig Stolarczyk, Sudarshan Pinglay, Sanjay Srivatsan, Carl de Boer, and Nicholas Mateyko for helpful discussions and insightful feedback.

## Supplementary Materials

Supplementary data are available online.

## Data Availability

Raw PacBio sequencing reads generated from assembled gene libraries were deposited in the NCBI Sequence Read Archive (SRA) and are publicly available under BioProject accession PRJNA1469590 (https://www.ncbi.nlm.nih.gov/bioproject/PRJNA1469590). The complete DropSynth-Gold oligonucleotide design pipeline used for fragmenting input gene sequences into Golden Gate-compatible oligonucleotide assemblies is publicly available on GitHub through the DropSynth-Gold repository (https://github.com/PlesaLab/DropSynthGold). The repository includes workflows and scripts for sequence preprocessing, codon optimization, restriction site filtering, oligonucleotide splitting, and generation of Golden Gate overhang-compatible oligo designs for multiplexed gene assembly.

