## Supplementary Materials for "DropSynth-Gold: Golden Gate Assembly in Emulsions Extends Multiplexed Gene Libraries to Greater Lengths"

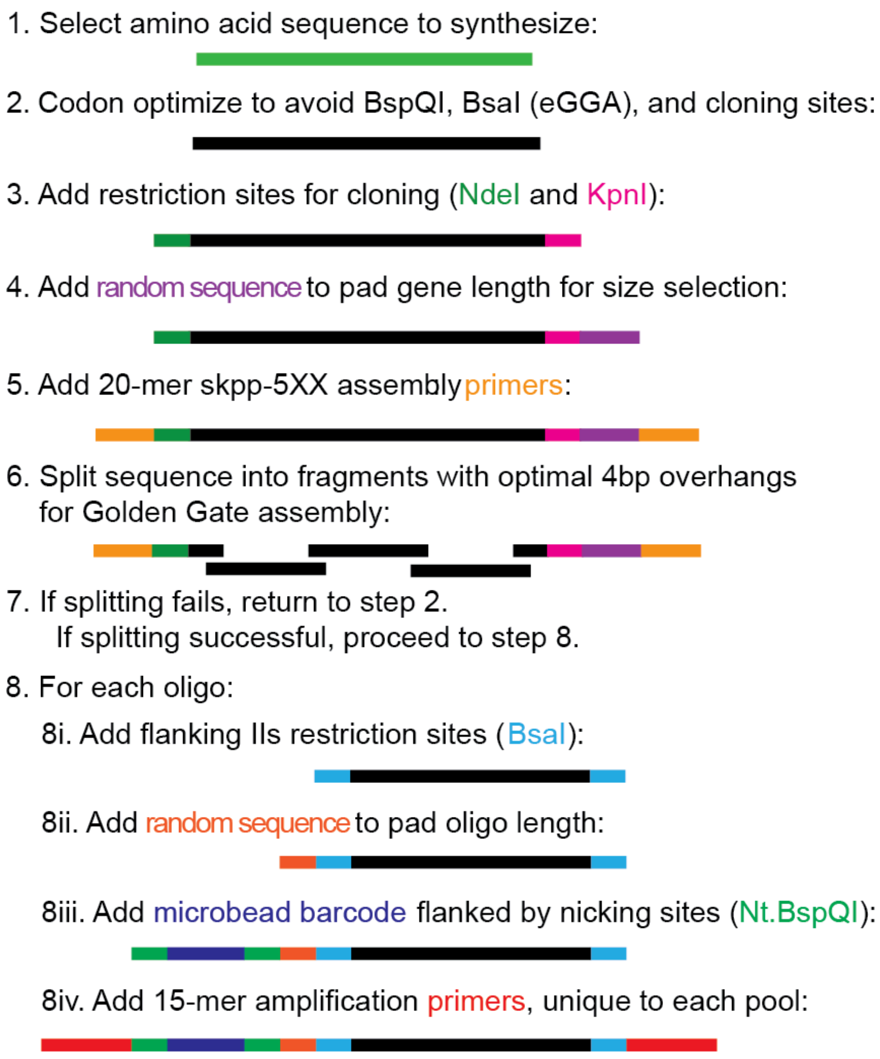


**Figure S1. Overview of the DropSynth-Gold oligo design workflow.** The DropSynth-Gold oligo design pipeline, available at https://github.com/PlesaLab/DropSynthGold, takes as input amino acid sequences and generates oligonucleotide libraries for Golden Gate-based emulsion assembly. Protein sequences are first codon optimized while removing forbidden restriction enzyme recognition sites, including BspQI and BsaI, as well as user-defined cloning sites. Cloning restriction sites (e.g., NdeI and KpnI) and 20-mer skpp-5XX assembly primer sequences are then added to the gene construct. To enable size selection and pooled processing, random padding sequence may be incorporated to normalize construct lengths across a library. The finalized construct is subsequently split into assembly fragments using optimized 4-bp Golden Gate overhangs selected to minimize secondary structure and maximize assembly compatibility. If fragment splitting fails because of unfavorable overlap properties, homopolymers, repetitive sequence content, or unresolved restriction sites, the sequence is reassigned alternative codons and the design process is repeated iteratively until a valid solution is identified. For each resulting oligonucleotide, flanking Type IIS restriction sites (BsaI) are added to enable Golden Gate assembly, followed by optional random padding to normalize oligo length. Oligos are then appended with a microbead barcode sequence flanked by nicking enzyme recognition sites (Nt.BspQI) to enable bead-linked assembly within emulsions. Finally, pool-specific 15-mer amplification primer sequences are added to support amplification and recovery of oligo libraries from the synthesized oligo pool.

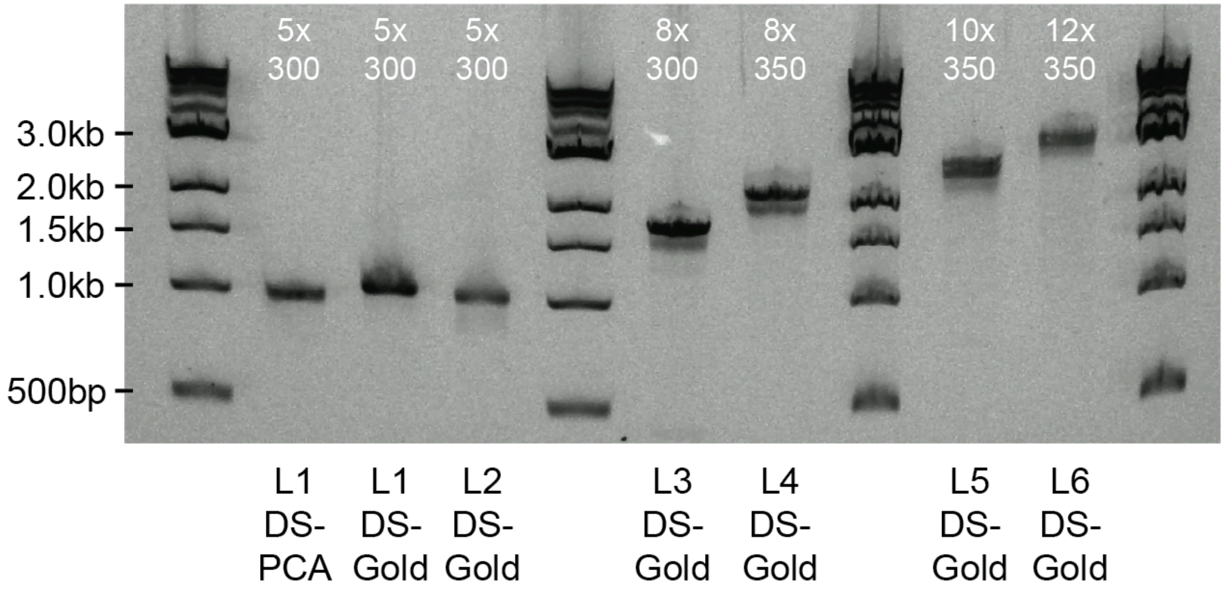


**Figure S2. Agarose gel analysis of assembled DropSynth and DropSynth-Gold libraries spanning increasing construct lengths and fragment counts.** Assembly products from PCA-based DropSynth (L1 DS-PCA) and DropSynth-Gold libraries (L1-L6) were analyzed on a 1% agarose E-Gel following pooled assembly and amplification. Libraries span assembly architectures ranging from 5x300-mer to 12x350-mer designs, corresponding to expected construct sizes from ~1.0 kb to ~3.0 kb. Lane identities are indicated below each sample, with assembly architectures shown above each lane. DNA ladders are shown for size reference.

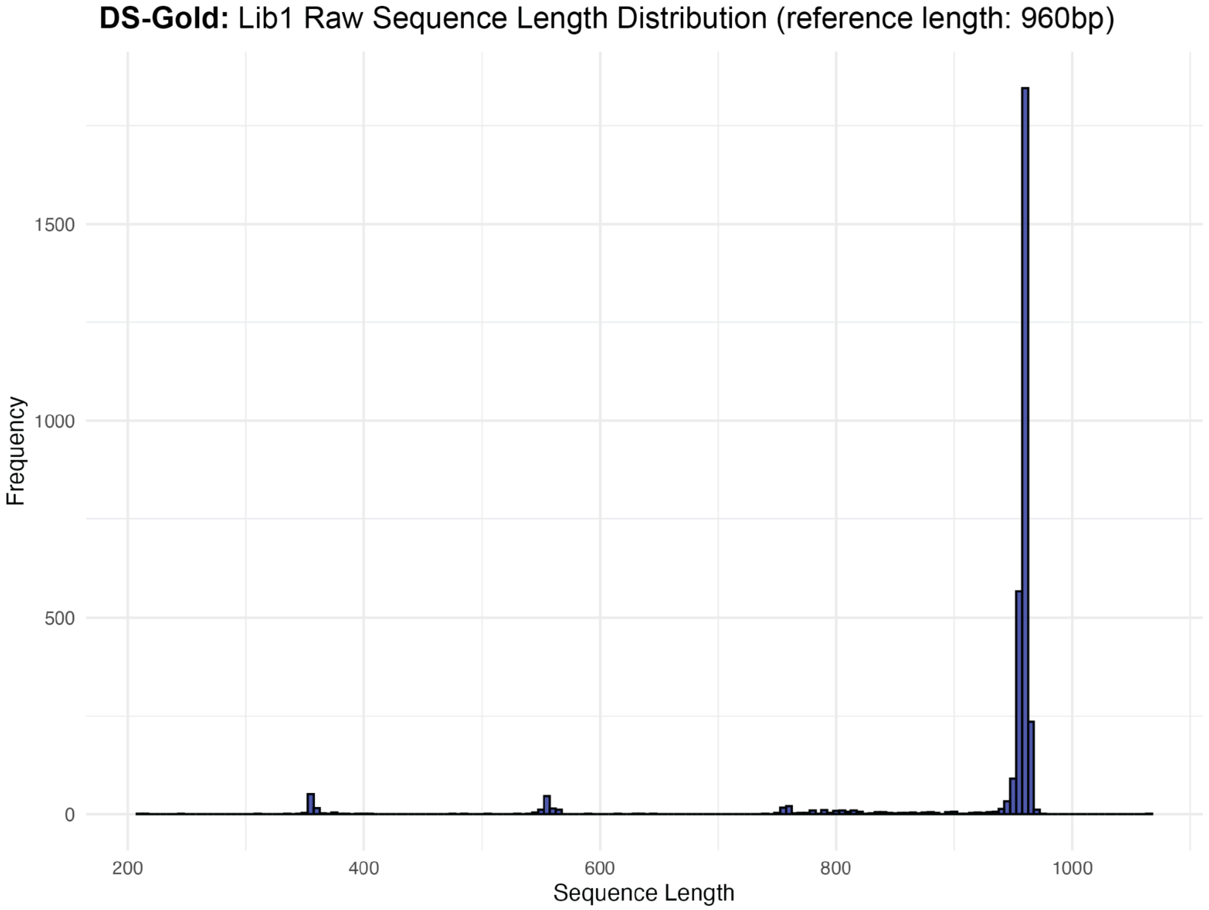


**Figure S3. Distribution of raw sequencing read lengths for DropSynth-Gold assemblies.** Histogram showing raw sequence read lengths for the DropSynth-Gold Lib1 assembly (5x300-mer design; expected reference length: 960 bp). A dominant and sharply defined peak corresponding to the full-length 5-fragment assembly was observed near the expected construct length. Smaller discrete peaks corresponding to shorter partial assemblies, including apparent 2-, 3-, and 4-fragment products, were present at substantially lower abundance. Unlike the broad smeared byproduct distributions commonly observed in PCA-based assemblies, the discrete nature of GGA intermediates produces well-separated length populations, enabling more effective size selection and enrichment of correctly assembled full-length products.
